# Virus transmission by ultrasonic scaler and its prevention by antiviral agent

**DOI:** 10.1101/2021.02.10.430630

**Authors:** Aleš Fidler, Andrej Steyer, Rok Gašperšič

## Abstract

During an ultrasonic scaler (USS) operation, droplets and aerosol are generated that may contribute to the transmission of viruses contained in saliva and gingival crevicular fluid. The purpose of this research was to develop an experimental model for testing the spread of viruses during USS instrumentation and to examining the prevention of spreading by replacing the coolant with an antiviral agent. In a virus transmission tunnel, USS operation with saline coolant and delivery of a viral suspension to the vicinity of USS tip generated droplets and aerosol containing Equine Arteritis Virus (EAV). Evaluation of droplet transmission was evaluated with adherent 48h cell culture monolayer RK13 cell lines in standard 48-well-plates positioned at a distance from 30 to 55 cm. The aerosol was collected by a cyclone aero-sampler flow of 100l/min. Antiviral activity of 0.25% sodium hypochlorite or electrolyzed water (EOW) was tested by suspension test. The two tested antiviral agents’ transmission prevention ability was evaluated by repeating the same experiment as with saline coolant. All experiments were repeated twice. With saline coolant, the cytopathic effect on cells was found in cells up to the distance of 45 cm, with the number of infected wells decreasing with distance. Viral particles were detected in only one AS in a very low concentration (≤4.2 TCID50/ml). In suspension test of 0.25% NaOCl and EOW, the TCID50/ml was below detection limit after 5s. With both antiviral agents, no cytopathic effect was found. However, the cytotoxic effect of 0.25% NaOCl was evident up to the distance of 35 cm. By USS activity, EAV could be transmitted by droplets up to a distance of 45 cm. Both antiviral agents could prevent virus droplet transmission. The transmission of EAV by aerosol yielded inconclusive results.

## Introduction

It is well recognized that dental procedures represent a potential way of infection transmission due to extensive generation of droplets and aerosol by using power-driven instruments, such as ultrasonic scaler (USS), high-speed rotary instruments, and air-and-water syringe. In the past, the spread of bacteria associated with dental procedures has already been extensively studied in real or artificial working environment settings, reporting on large contamination to operator’s face and head, patient chest and also surfaces up to 3 meters from the patient mouth (Innes et al. 2021). To reduce bacterial load, preprocedural mouth-rinse was suggested despite the moderate level of evidence to support such action (Marui et al. 2019). The bacterial content of aerosols was reduced 2 to 7 fold by using povidone-iodine and chlorhexidine gluconate as an ultrasonic liquid compared with distilled water in an *in-vivo* study (Jawade et al. 2016).

With the COVID-19 pandemic, the focus of dental procedure associated transmission has rapidly changed from bacteria to viruses. Although the SARS-CoV issue in 2004 was recognized in dental literature (Li et al. 2004a; Li et al. 2004b), no research is presently available on virus transmissions associated with dental procedures (Koletsi et al. 2020). It is generally acknowledged that dental procedures represent a high risk of SARS CoV-2 virus transmission(Ather et al. 2020; Meng et al. 2020; Sabino-Silva et al. 2020; Izzetti et al. 2020) due to its presence in saliva (To et al. 2020), gingival crevicular fluid (Gupta et al. 2020), nasopharyngeal, oropharyngeal and bronchial excretions (Liu et al. 2020). For the provision of safe dental care for dental team members and patients, numerous preventive measures were proposed and included in professional (Ather et al. 2020; Meng et al. 2020; Sabino-Silva et al. 2020; Izzetti et al. 2020) and national (Clarkson et al. 2020) COVID-19 prevention guidelines. The measures range from pre-procedure mouth-rinse, use of personal protective equipment, high volume evacuation, air purification (ventilation, filtering, air disinfection), and surface disinfection. Personal protective equipment can protect the dental care providers; however, the contamination of clinical environments by sprays still necessitates periods of “fallow time” between appointments to protect patients and staff (Sergis et al. 2020). A special concern was raised for open plan clinic environments, reporting a safe distance of 5m (Holliday et al. 2021) Although preprocedural mouth-rinse is included in most recommendations (Clarkson et al. 2020), its effectiveness is questionable as no scientific clinical evidence exists for reducing the viral load (Carrouel et al. 2020). So far, antiseptic mouth rinses containing cetylpyridinium chloride or povidone-iodine have shown the highest potential to reduce viral load in infected subjects (Herrera et al. 2020). Thus, there is a high need for effective prevention measures against virus transmission in dentistry. The splatter and aerosol source is a mixture composed of virus-containing body fluids and water from the dental unit water system.

Although a simple replacement of cooling liquid with virucidal agent might have a potential to reduce viral spread, this possibility has not been evaluated. Such an approach would require a highly effective agent capable of virus inactivation in a very short time. Several regents were already found to be effective against coronaviruses in as short as 30 s of suspension test (Kampf et al. 2020). Some of them, like hydrogen peroxide (Boyd 1989) or sodium hypochlorite (Rich and Slots 2015) have already been evaluated for safe use as an oral rinse. Additionally, the electrolyzed water was proposed as an effective and safe biocidal agent for nasal (Yu et al. 2011) or ocular (Sipahi et al. 2019) application.

The aim of the study was twofold, (i) to develop an experimental model for evaluation of viral transmission by USS activity, and (ii) to evaluate reduction of viral transmission with USS by the replacement of water with a virucidal agent.

## Methods

To evaluate our hypothesis, we have designed a virus transmission tunnel (VTT), thus simulating dental procedure in controlled laboratory conditions. It featured the operating USS in the presence of virus suspension and virus sampling device for a spill, droplets, and aerosol. It consisted of a clear acrylic box, resembling a similar device (Singanayagam et al. 2020), with dimensions of 120 x 50 x 40 (L x W x H) manufactured specifically for this purpose (Acrytech, Ljubljana, Slovenia).

At one end of the VTT, a platform for droplet and aerosol generation was positioned 25 cm above the floor. The piezoelectric USS (Varios 970, NSK dental, Nakanishi Inc. Shimohinata, Japan) with the handpiece (VA2-LUX-HP) and a general prophylactic tip (G4) was firmly positioned in a mechanical holder. For all experiments, the USS was set to General mode, power level to 10 and the coolant level to 3, and flow rate of 17.5 ml/min. The USS tip was positioned into a semi-cylindrical groove, measuring 30 x 3 x 3 mm (LxWxD) (Fig 1). In the same groove, a virus suspension was delivered by an IV administration set (Normal set, Ferrari L., Verona, Italy)with a flow rate of 104 ml/h (1,73ml/min) through a pre-curved, 20G blunt, rounded tip needle (0.9X 42mm) (Transcodent, Kiel, Germany), stably mounted with a connecting element (Fig. 1).

**Figure 1.**
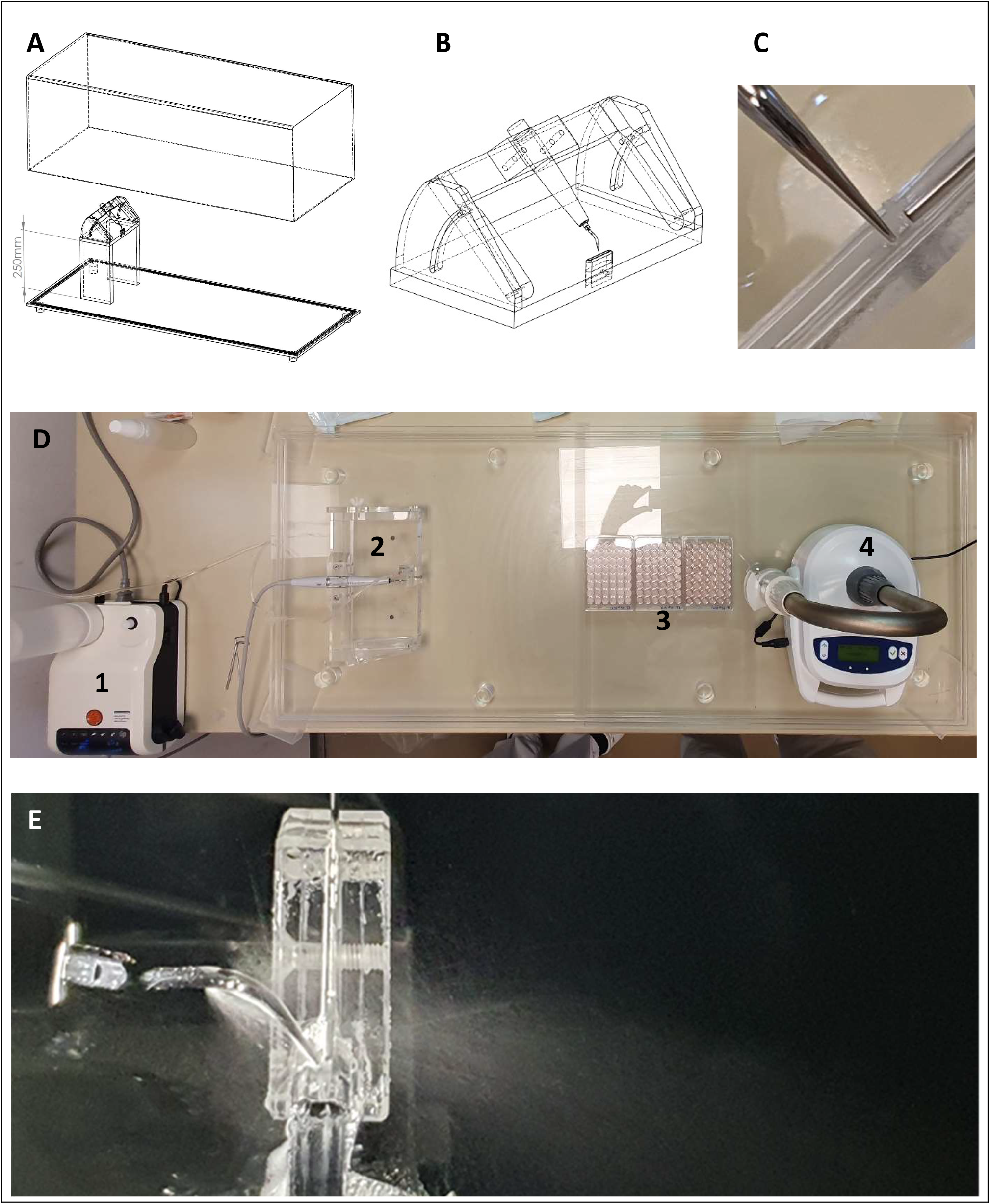
Experimental set-up for virus transmission evaluation. **(A)** - The schematic diagram of the virus transmission tunnel (VTT). **(B)** – Platform with ultrasonic scaler (USS) handpiece holder and groove, **(C)** – semi-cylindrical groove with USS tip, and blunt needle for virus suspension delivery, **(D)** Top view on the virus transmission tunnel, showing **(1)** - USS device, **(2)**– platform with USS handpiece and groove, **(3)** - three consecutively positioned 48-well cell culture plates, and **(4)**- Air Sampler. **(E)** - USS tip, positioned in the groove, during experiment.

### Virus suspension and sampling

For experiments, a laboratory Equine Arteritis Virus (EAV) strain was used. EAV was propagated in a 48 h cell culture monolayer. A specific cytopathic effect (CPE) with cell rounding and degeneration of cell monolayer is observed in 48h post-inoculation in 75 cm^2^ cell culture flasks. After CPE reaches 90% of the monolayer, the cell culture flask was freeze-thawed two times, and finally, the whole-cell debris and media were transferred to 50 ml centrifugation tubes. Cell debris was removed by centrifugation 15 min at 1500×*g*. Clear virus suspension was stored at -80°C in 1 ml cryotubes. One ml of clear virus suspension with a concentration of 1.33×10^6^ TCID_50_/ml (infective virus units: 50% Tissue Culture Infectious Dose per milliliter) was inoculated into 250 ml sterile saline solution (0,9% NaCl), which was delivered to the position of USS tip through infusion system. Virus transmission in the VTT was achieved by simultaneous activation of USS and flow of virus suspension in a duration of 10 min.

#### Virus sampling

Virus sampling was performed at three sites: liquid sample (LS), droplet sample (DS), and aerosol sample (AS). At the end of the procedure, the LS was acquired from a mixture of virus suspension and cooling liquid, flowing freely from the groove and collected in a glass jar placed directly under the platform. The DSs were acquired during the procedure by the three 48-well standard cell culture plates (8.5 × 12.8 cm) (Cell Culture Multiwell Plate, Cellstar) with adherent 48h cell culture monolayer that were positioned consecutively from the distance of 30 cm to 55 cm. In total, 108 wells, arranged in 18 lines, with six wells per line, were used as DS. After the procedure, plates were covered, removed, and incubated. The AS was performed using a cyclone air sampler (Coriolis Micro, Bertin Technologies, France), with airflow set at 100 l/min (Fig 1X). The air inlet was positioned at a distance of 60 cm from the USS tip and height of 25 cm. The experiment was repeated twice.

### Virus transmission evaluation

The virus was efficiently propagated in an adherent RK13 cell line (source: *Oryctolagus cuniculus*, kidney, epithelial). Eagle’s Minimum Essential Medium (Sigma-Aldrich, MO) was used for cell culture medium, supplemented with final 10% fetal bovine serum (EuroClone, Italy) concentration.

Virus concentration in LS and AS was determined with titration on cell monolayer in 96-well microtiter plate. Each 10-fold dilution was inoculated in 8 replicates, including negative control. Briefly, cell culture media was removed from the cell monolayer, and 100 µl of virus suspension (LS or AS sample) was added to each well. The inoculated microtiter plate was incubated for up to 4 days in 37°C, 5% CO_2_ atmosphere, and 95% humidity. After four days, cells were screened for CPE, and the number of infected wells was recorded.

Virus transmission in DS was evaluated by CPE formation in exposed microliter plates incubated for up to 4 days in 37°C, 5% CO_2_ atmosphere, and 95% humidity. After four days, cells were screened for CPE, and the number of inoculated wells in each row was recorded.

### Irrigant suspension evaluation

The virucidal ability of two irrigants was evaluated with a suspension test. A 0.25% NaOCl and EOW (Table 1) were used. The pH, redox potential (Multi 3630 IDS, WTW, Weilheim, Germany), and free chlorine (Pocket Colorimeter II, Loveland, Colorado, USA) were measured and recorded (Table 1). Virus suspension was exposed to the agent, and a reduction of infective virus particles was evaluated after a contact time of 5s, 30s, 1 min, 5 min, and 10 min.

**Table 1.**
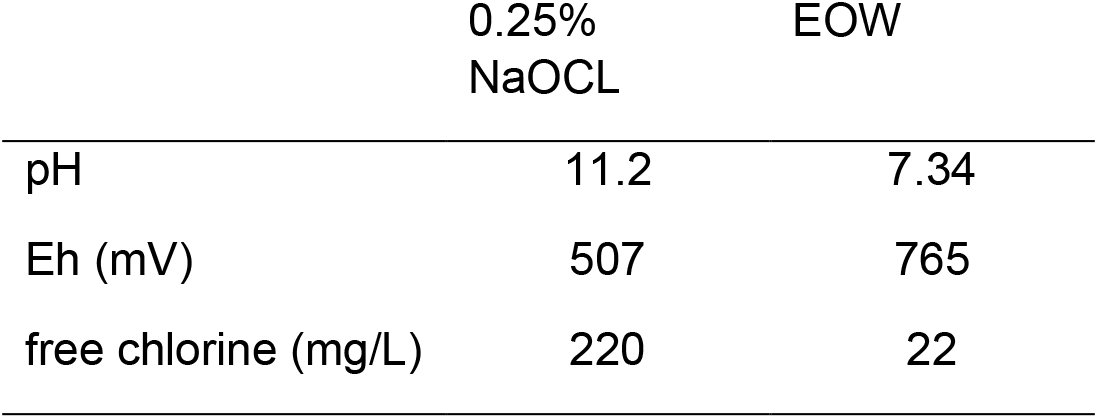
Chemical characteristics of antiviral coolants, used in suspension test and virus transmission prevention test

### Virus transmission prevention evaluation

The virus transmission prevention test was evaluated with the same experimental setup as for the virus transmission test, except that sterile saline as a cooling agent was replaced with agents evaluated in suspension tests: 0.25% NaOCl in the first and EOW in the second experiment. The irrigant flow rate was the same as in the virus transmission test. For each irrigant, the experiment was repeated twice.

## Results

### Virus transmission test

The estimated virus concentration in the LS sample, collected in the vicinity of the droplet-generating platform, was 3.25 log TCID_50_/ml. In DSs collected in wells, CPE associated with viral infection was found in the first 16 of 18 lines of cultivation plates. More than one infected well per line was found only in the first seven lines (Fig. 2A). In one of two AS, viral particles were detected in a very low concentration, which was ≤0.63 log TCID_50_/ml.

**Figure 2.**
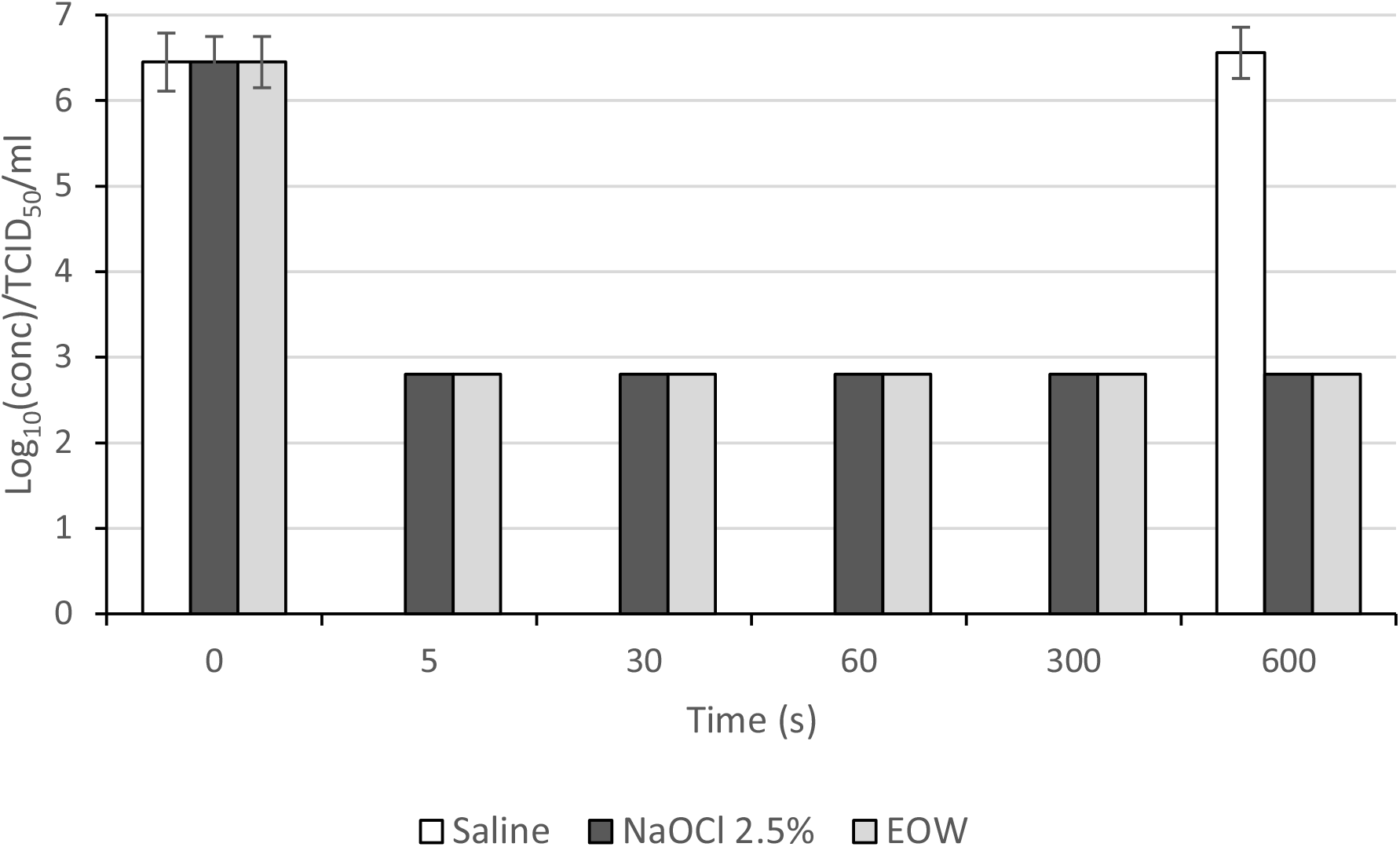
Suspension test for the saline, 0.25% NaOCl, and electrolyzed water.

### Suspension test

The suspension test showed a highly efficient reduction of virus concentration in a short time (Fig 3). In 5 second contact time, both the EOW and the 0,25% NaOCl decreased the log TCID50/ml concentration from 6.45±0.60 to a value of ≤2.8, which was the limit of detection in our test due to virus dilution. The log_10_ (conc)/TCID_50_/ml value remained unchanged after more extended periods of observation. The suspension test with saline did not affect the virus concentrations (Fig 3).

**Figure 3.**
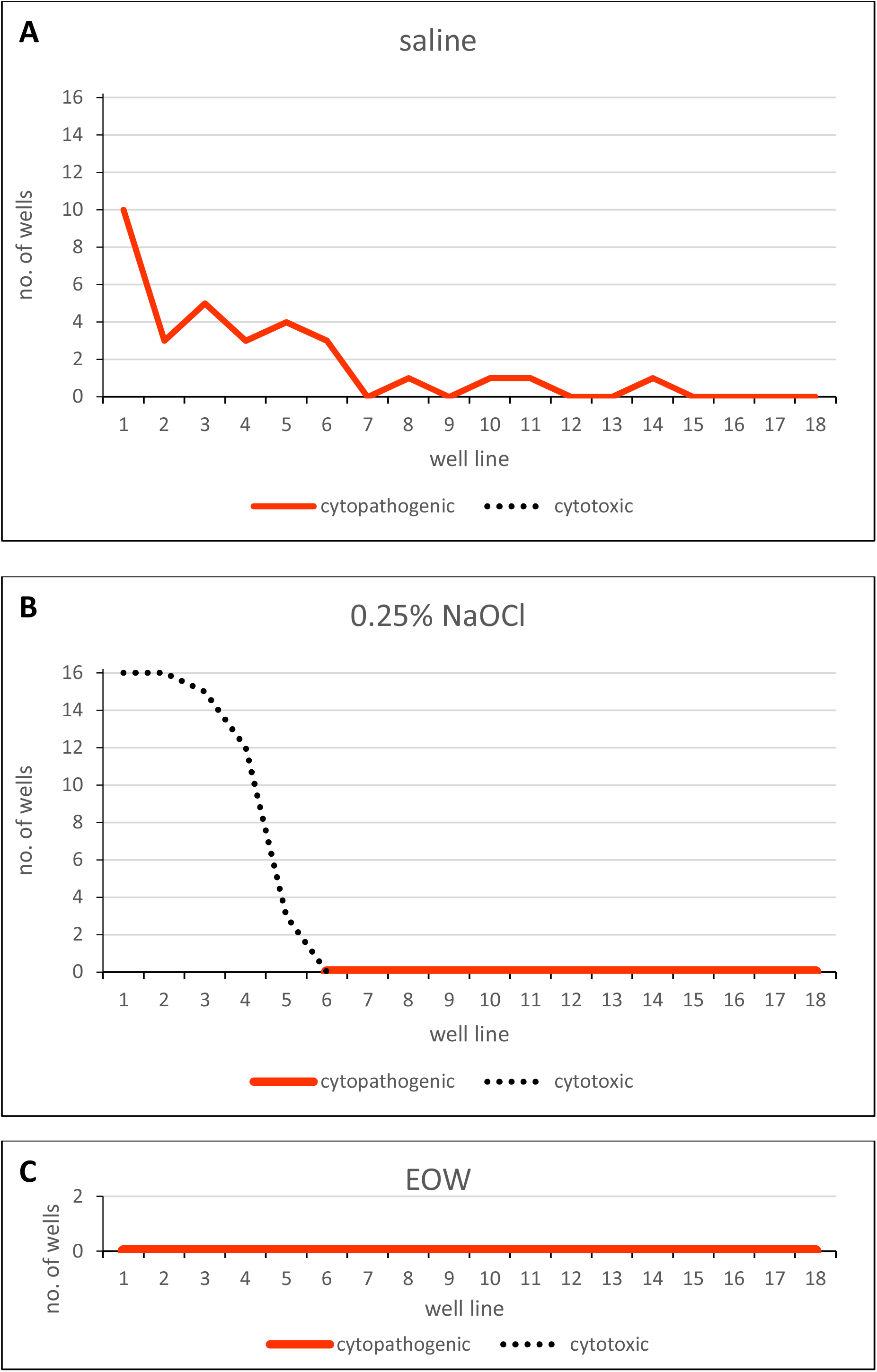
The cytopathogenic and cytotoxic effect in droplet samples, obtained from three different coolants. **(A)** saline, **(B)** 0.25% NaOCl, and **(C)** electrolyzed water. In the case of cytotoxic effect, the cytopathogenic could not have been evaluated.

### Virus transmission prevention test

In LS, for both irrigation experiments, no infective viral particles were detected. Importantly, a cytotoxic effect of NaOCl was found in LS for dilutions up to 10^5^, while no cytotoxic effect was found for EOW.

In DS, sampled with 0.25% NaOCl as an irrigant (Fig 2B), the cytotoxic effect was found in the first five lines, and thus the effect of infective viruses (CPE) could not be evaluated. In lines from 6 to 18, no CPE nor cytotoxic effect was found. With EOW as a coolant (Fig 2C), nor CPE neither cytotoxic effect was found in any line.

In AS, for both irrigation experiments, no infective viral particles were detected. However, for 0.25% NaOCl cytotoxic effect was found at 10^−1^ dilution.

## DISCUSSION

The study results indicate that EAV was transmitted mainly by droplets and up to the distance of 45 cm by USS activity. The transmission of EAV by aerosol, however, yielded inconclusive results. More importantly, the virus droplet transmission could be prevented by a simple saline replacement with a virucidal agent that has already been tested as mouth-rinse for home oral care.

Our aim was to select a non-human viral pathogen, which resembles a similar structure and genome as SARS CoV-2. An Equine Arteritis Virus (EAV) is an animal pathogen, which is species-restricted to *Equidae* members. EAV is an enveloped virus with a single-stranded, positive-sense RNA genome. EAV is also a member of Baltimore’s IV group and presents a similar structure to Coronaviruses, similar stability in the environment, and inactivation by general disinfectant as SARS CoV-2 (Balasuriya et al. 2018; Chan et al. 2020). Besides, EAV is also transmitted through respiratory route and aerosol transmission, although indirect or close contact is required for infection (Balasuriya et al. 2018).

In the first part of the study, we designed an experimental setup to demonstrate the EAV spread by USS action. The EAV predominantly spreads via larger droplets (splatter) that, according to ballistic laws, reach the near surrounding area, as infected droplets consistently infected cells on plates up to the distance of 45 cm (16^th^ well) from the USS tip. These observations are in accordance with a recent review (Sergis et al. 2020) stating that contaminants settle to a great extent on the operator’s dominant arm, eyewear, and chest of the patient, and to a lesser extent on the non-dominant arm and chest of the operator and assistant. To the best of our knowledge, this is the first proof of the contribution of an USS action to the spread of the virus via droplets to the environment. Although we were able to detect EAV in low concentration in air sample, we failed to achieve the reproducible collection of the infectious virus from the air-sampler. This observation is in accordance with a recent review on air contamination by SARS-CoV-2 in hospital settings (Birgand et al. 2021). Air samples from the close patient environment were positive for SARS-CoV-2 RNA and viable viruses only in 17.4 % and 8.6%, respectively. As the volume of a typical aerosol particle with a diameter of 1µm is 5.23×10^−13^ ml, and 3.58×10^11^ of such particles may be generated from 1 ml, it is possible to calculate by dividing the number of virus particles in 1 ml by the number of aerosol particles, obtained from 1 ml suspension of our study. i.e. 1.78×10^3^ / 3,58×10^11=^ 4.97×10^−9^, representing a fraction of particles actually harboring a virus. This might be the explanation for low percentage of aerosol virus transmission. Another reason could be the virion damage during the sampling process (Pan et al. 2019).

In our experiments, the clear virus suspension with a concentration of log 4.12±0.60 TCID_50_/ml and flow of 1.73 ml/min was diluted with irrigant with flow between 12.5 and 17.5 ml/min. The concentration of log 3.25±0.59 TCID_50_/ml was measured in LS. This observation is in accordance with a recent paper, stating that mixing the introduced coolant with real saliva also requires consideration (Sergis et al. 2020). In our case, with virus titration control in saline suspension and splatter collected directly at the USS, the concentration reduction was 0.87 log, which is near 10x dilution and goes perfectly with the ratio of virus suspension and irrigant flow. This observation confirms the importance of dilution, stated in previous studies (Epstein et al. 2020; Holliday et al. 2021)

In the second part of the experiment, we managed to prevent the droplet spread of the virus to the surroundings by changing the saline coolant with EOW or hypochlorite. Despite the promising results of our preliminary tests, in which 0.25% hypochlorite did not show considerable cytotoxicity, such effect was always found in the wells reached by a higher number of droplets (the first six lines). Nevertheless, no cytotoxic effect was found in the wells at a larger distance from the USS tip. It should be noted that in the experiment with the saline, cell infection was apparent to the much greater distance (up to the 16^th^ well). Therefore, the absence of CPE in wells 6-16 can be attributed to the effective inactivation of the virus by 0.25% hypochlorite.

However, the effective infection prevention of cells was particularly interesting for the EOW experiment, as, despite the absence of cytotoxicity, we never found a single well with CPE suggestive of viral infection. Since all experiments were well controlled and performed sequentially, it is unlikely that the absence of signs of viral infection in these experiments resulted from a methodological error. We may consider our experiments as the first that have clearly shown the possibility of disinfecting the virus in droplets generated by USS by replacing inert coolant with antiseptic irrigant. In addition, EOW showed a high virucidal effect in the suspension test. A low level of cytotoxicity and the high virucidal effect are optimal for highly effective potential disinfectants in dental procedures. Considering the fact that AS from the NaOCl experiment exhibited cytotoxic effect already at 10^−1^ dilution, we could assume that NaOCl-containing aerosol was successfully collected during the experiment. This result could be attributed to the high throughput air sampler used in our case, as in 10 minutes, the whole volume of the VTT was filtered 4-times. Nonetheless, the detection of infective virus particles was not successful in the first part of the experiment.

Besides direct infection by droplets, the proposed approach might also effectively prevent indirect transmission by contaminated surfaces or reduce viral load. It has been demonstrated that SARS CoV-2 might be transmitted indirectly via contaminated surfaces as the virus can survive on surfaces like metal, glass, or plastic for up to a couple of days (Otter et al. 2016; Kampf et al. 2020).

To the best of our knowledge, this is the first dental procedure virus transmission study, performed with viable virus evaluation. The proposed setup also allows evaluating other powered dental devices, such as air turbines, high-speed contra-angles, and dental lasers. As EAV is incapable of infecting humans, the experiments can be safely performed using standard PPE with no risk for the researchers. However, the combination of EAV and RK13 cell line, representing kidney epithelial cells, does not precisely reflect clinical conditions, featuring SARS CoV-2 and respiratory epithelial cells. The use of SARS CoV-2 would require facilities with Biosafety Level 3 conditions, and with inducing aerosols, the whole experiment procedure would not be possible.

EOW was found to have no systemic effects when it was provided as by drinking water to mice during two weeks experiment, which served to the authors as indirect proof for safe usage as a mouthwash (Morita et al. 2011). EOW was already proposed as an effective and safe biocidal agent for nasal (Yu et al. 2011) or ocular (Sipahi et al. 2019) application. Similarly, the 0.25% sodium hypochlorite has been proposed for oral rinse (Galván et al. 2014; Gonzalez et al. 2015).

It should be noted that our experiments were performed without any splatter or aerosol reduction device in order to maximize the contamination and to test the effect of coolant in the most challenging situation. Using a high vacuum evacuator or similar device, as they are usually used in a clinical setting, would considerably reduce the spread of splatter and aerosol (King et al. 1997; Holloman et al. 2015). Similar to a recent study on atomization from rotary dental instruments (Sergis et al. 2020), the generalization of our findings into the wide variety of clinical settings is difficult due to variability of USS devices and their operating parameters, including power setting tip selection and irrigant flow rate. Even larger differences may be found between dental procedures (Allison et al. 2021).

In conclusion, by using the proposed experimental model, we managed to predictably demonstrate virus spread by droplets due to USS action in virus suspension. More importantly, we managed to mitigate the virus spread by a simple substitution of the irrigant with already clinically used virucidal agents, namely sodium hypochlorite and EOW. Future research using the proposed experimental model should include other aerosol and droplet generating dental procedures, additional agents, and ultimately the SARS-Cov-19 virus before clinical application. Nevertheless, the proposed principles for virus spread mitigation seems promising and warrant further evaluation.

## Acknowledgments

Laboratory Equine Arteritis Virus (EAV) strain was kindly provided by the Institute of Microbiology and Parasitology, Veterinary Faculty, University of Ljubljana, Slovenia.

The work was supported by the Ministry of Higher Education, Science, and Technology, Republic of Slovenia, under grant number P3-0293 and P3-0083.

## Notes

### Competing Interest Statement

The authors have declared no competing interest.

### Summary of Updates

No revision needed. Only the title page was added.

